# Mast cell function in prostate inflammation, fibrosis, and smooth muscle cell dysfunction

**DOI:** 10.1101/2021.03.23.436678

**Authors:** Goutham Pattabiraman, Ashlee J. Bell-Cohn, Stephen F. Murphy, Daniel J. Mazur, Anthony J. Schaeffer, Praveen Thumbikat

## Abstract

Intraurethral inoculation of mice with uropathogenic *E. coli* (CP1) results in prostate inflammation, fibrosis, and urinary dysfunction, recapitulating some but not all of the pathognomonic clinical features associated with benign prostatic hyperplasia (BPH) and lower urinary tract symptoms (LUTS). In both patients with LUTS and in CP1-infected mice, we observed increased numbers and activation of mast cells and elevated levels of prostate fibrosis. Therapeutic inhibition of mast cells and the histamine 1 receptor in the mouse model resulted in reduced mast cell activation in the prostate and significant alleviation of urinary dysfunction. Treated mice showed reduced prostate fibrosis, less infiltration of immune cells, and decreased inflammation. In addition, as opposed to symptomatic CP1-infected mice, treated mice showed reduced myosin light chain (MLC)-2 phosphorylation, a marker of prostate smooth muscle contraction. These results show that mast cells play a critical role in the pathophysiology of urinary dysfunction and may be an important therapeutic target for men with BPH/LUTS.

**NEW AND NOTEWORTHY:** LUTS-associated BPH is derived from a combination of immune activation, extracellular matrix remodeling, hyperplasia, and smooth muscle cell contraction in prostates of men. Using a mouse model, we describe the importance of mast cells in regulating these multiple facets involved in the pathophysiology of LUTS. Mast cell inhibition alleviates both pathology and urinary dysfunction in this model suggesting the potential for mast cell inhibition as a therapeutic that prevents and reverses pathology and associated symptomology.

## INTRODUCTION

Lower urinary tract symptoms (LUTS) occur at high incidence in both men and women with increasing age. Benign prostatic hyperplasia (BPH) associated LUTS is a major contributing factor to negative quality of life in men, costing over $1 billion dollars in healthcare costs, in the United States alone, per year as of 2014 (1). In men, LUTS is multifactorial but is commonly associated with the development and progression of BPH. BPH is a histological condition that is characterized by varying combinations of epithelial and stromal hyperplasia within the prostate, and is prevalent in over 50% men of age over 50 (2, 3).

Historically, risk factors linked to the development and progression of LUTS include age, genetics, infections, stress, and BPH (4, 5). Studies have revealed prostate inflammation to be highly associated with LUTS, correlated with prostatic enlargement, and implicated as a cause of prostatic fibrosis (6). Tissue inflammation and fibrosis has numerous potential causes including aging, infection (from bacterial prostatitis or urinary tract infections), dietary habits, hormonal changes and physical trauma and urinary reflux (5, 7–10). Inflammation and fibrosis are thought to cause increased rigidity of the prostate, resulting in increasing pressure and constriction on the urethra, leading to LUTS (5, 6, 11, 12). Alongside inflammation and fibrosis, aberrant smooth muscle contraction also causes voiding symptoms in patients by impairing the bladder outlet due to urethral obstruction, a condition commonly referred to as bladder outlet obstruction (BOO) (13).

Mast cells act as crucial cellular sensors and regulators of inflammation, fibrosis, and smooth muscle cell contraction (14–16). In addition to playing a critical role in allergic immunity, mast cells can sense various “danger signals” by using the plethora of pattern recognition receptors expressed on both the cell surface and within the cells. Activation of these receptors induces release of a milieu of cytokines which influence immune cell infiltration and activation, and release of various other factors including chymases, tryptases and proteases which can interact with and alter the local tissue environment leading to tissue fibrosis, repair and remodeling (14–21). Clinical reports suggest that mast cells play an important role in the development and persistence of inflammation in BPH associated LUTS (22, 23). Mast cell secreted factors, predominantly in the context of airway smooth muscle cells, have been implicated to play an important role in controlling smooth muscle cell growth, differentiation, and contraction (24–26). Also, we have previously shown that protease activated receptor 2 (PAR2) activation by mast cell derived proteases in prostate smooth muscle cells causes cellular contraction (27). Given the evidence that prostate fibrosis and inflammation and smooth muscle cell contraction are important in the development of LUTS, and the influence that mast cells have on such processes, we assessed the role of mast cells in the development of LUTS.

We have previously reported the ability of an uropathogenic clinical prostate isolate of *E. coli* (CP1) to colonize the prostates of mice and recapitulate aspects of LUTS such as urinary dysfunction, prostate inflammation, and fibrosis. These changes and observations persist long after bacterial clearance (28, 29). In this study, we observed the presence of an increased number and activation of mast cells in the prostate of CP1 infected mice that was associated with increased inflammation and the development of fibrosis. Upon inhibition of mast cell activity using a combination a mast cell stabilizer (MCS) and histamine 1 receptor antagonist (H1RA), findings of urinary dysfunction were ameliorated and pathological features such as immune cell infiltration, fibrosis and smooth muscle contraction were significantly reversed. These studies implicate mast cells as critical mediators of prostate pathology in LUTS and furthermore show that a novel and innovative treatment approach that blocks release of mast cell factors and modulates the immune response may be useful in alleviating LUTS. This may be a promising novel therapy to prevent or more effectively treat LUTS in aging men.

## MATERIALS AND METHODS

### Mice

Male C57BL/6 mice (5-7 weeks of age) were obtained from Jackson Laboratory. Mice were housed in a single environmentally controlled room within the Northwestern University animal facility. All animal experiments and procedures have been approved by the Northwestern University Animal Care and Use Committee.

### Trans-urethral mice infections

Mouse infections were performed as described previously (28–30). Briefly, CP1 *E. coli* bacteria were grown in LB overnight shaking at 37°C, followed by overnight static subculture at 37°C. The next day, bacteria were concentrated at 2 × 10^10^ bacterial/mL in PBS, and 10 μL (2 × 10^8^ bacteria) was instilled transurethrally into isoflurane-anesthetized male C57BL/6 mice. Age-matched control male C57BL/6 animals received a transurethral (t.u.) instillation of PBS (Gibco, Paisley, UK) and were kept in separate cages.

### Voiding behavior testing

To assess the voiding behavior of mice, we utilized the Urovoid system (Med Associates, Fairfax, VT, USA), a noninvasive means of measuring voiding function using a modified procedure previously described (31). The Urovoid system is designed to assess conscious urinary voiding behavior (frequency and voiding volume) in unrestrained mice for prolonged periods without the need for surgery or catheter implantation. Briefly, mice are singly housed in chambers access to water for 5 hours. Urine was collected below the grated cage on a balance [mice feces is separated using a mesh above the balance], and urine weight was recorded over time (1-hour post housing the mice to allow for acclimatization). After the completion of these measurements, animals were returned to their home cages for future experimentation or euthanized as per experimental design. The data collected is then using the Urovoid voiding frequency analysis system (Med Associates, Fairfax, VT, USA). A representative graph from the analysis software is shown in Fig. 2A.

### Mouse tissue preparation

The prostates were harvested from mice after euthanizing as described previously (32). Depending on the experimental design, each prostate sample (separated by lobes) were either fixed in 10% formalin or frozen @ −80°C in TRIzol^TM^ reagent (Life Technologies Corporation, Grand Island, NY, USA), or frozen @ −80°C in 1X RIPA (Santa Cruz Biotechnology, Dallas, TX, USA) containing complete EDTA protease inhibitor cocktail and phosSTOP phosphatase inhibitor (Millipore-Sigma, Burlington, MA, USA), or processed for tissue digestion for flow cytometry as described below. Formalin samples were further processed and embedded in paraffin by the Northwestern University Mouse Histology and Phenotyping Core. The formalin-fixed paraffin-embedded (FFPE) samples were then sectioned (5 μm sections) and mounted on glass slides for staining. The Northwestern University Mouse Histology and Phenotyping Core performed H&E staining, as well as IHC for mast cell tryptase.

### *In vivo* administration of mast cell stabilizer and H1 receptor inhibitors

Male C57BL/6 mice were intra-peritoneally treated starting at either 5- or 25-days’ post-infection [“prophylaxis” or “treatment” groups respectively] with a combination of cetirizine di-hydrochloride at 2.5 mg/kg (H1 receptor inhibitor) and cromolyn sodium salt at 0.5 mg/kg (mast cell stabilizer) (Sigma, St. Louis, MO, USA) daily for 10 days as illustrated in (Fig. 4A). The doses for these compounds were based on previous studies from our group (33). Following voiding behavior testing on days 14 or 35 [“early treatment” or “late treatment” groups respectively], mice were euthanized as per protocol approved by the Northwestern University Animal Care and Use Committee, and tissues were prepared as described above.

### Histological and Immunohistochemical Assays

All histological and IHC staining and assays were performed on the anterior, ventral, and dorsolateral separately. 5 μm sections were processed for H&E staining (performed by the Northwestern University Mouse Histology and Phenotyping Core). Briefly, H&E staining on FFPE tissues were performed on a fully automated platform (Leica Autostainer XL; Lecia Biosystems, Buffalo Grove, IL, USA) using Harris Hematoxylin (Fisher Scientific, Hampton, NH, USA) and Eosin secondary counter stain (Lecia Biosystems, Buffalo Grove, IL, USA). Prostate inflammation was assessed using the classification system as described previously (34). Inflammation scoring was quantitated as follows: 0 - no inflammation, 1 - mild inflammation, 2 - moderate inflammation, and 3 - severe inflammation. For assessing extracellular collagen deposition picrosirius staining was performed as previously described (28). Briefly, 5 μm prostate sections on glass slides were rehydrated, and slides were stained in picrosirius red solution (Direct Red 80 and saturated picric acid, Sigma, St. Louis, MO, USA) for 16 hours. The sections were washed in two changes of acidified water, dehydrated in ethanol, cleared in xylene, and mounted with Krystalon (EMD Millipore, MA, USA). Stained sections were quantitated using NIH Image J software and percentage of collagen deposition were calculated. For detecting mast cells, we use toluidine blue staining, a metachromatic stain should stain mast cells red-purple (metachromatic staining) and the background blue (orthochromatic staining). Briefly, 5 μm prostate sections on glass slides were rehydrated, and sections were stained using 0.1% toluidine blue for 2-3 minutes, washed thrice using distilled water, dehydrated in ethanol, cleared in xylene, and mounted with Krystalon (EMD Millipore, MA, USA). Stained cells were counted in a blinded manner to quantitate levels of resting and activated mast cells.

Bright-field images and circularly polarized images were taken on a Leica DMLA microscope (Leica Microsystems Inc., Buffalo Grove, IL, USA) using a QImaging MicroPublisher 3.3 RTV camera (Teledyne Photometrics, Tucson, AZ, USA) and analyzed on Micro-manager, an open-source microscopy software (35).

### Human Tissue Microarray

The tissue arrays of Human prostate normal as well as hyperplasia punch biopsy sections (PR632) were purchased from US Biomax, Inc. (Derwood, MD, USA). Picrosirius staining for the sections were performed as described above. IHC staining of mast cell tryptase was carried using mouse anti-human Mast Cell Tryptase Antibody (catalog # 369402, RRID:AB_2566541, BioLegend®, San Diego, CA, USA) [performed by the Northwestern University Mouse Histology and Phenotyping Core]. Briefly, FFPE tissue slides were deparaffinized on an automated platform (Leica Autostainer XL; Lecia Biosystems, Buffalo Grove, IL, USA). Slides are treated with an antigen retrieval step using a sodium citrate solution at pH 6, in a pressure cooker. Sections were incubated overnight at 4°C with human mast cell tryptase antibody in a humid chamber. All IHC staining was completed using chromogenic enzyme substrate reactions with DAB (Agilent Technologies, Santa Clara, CA, USA). Secondary antibody incubation and chromogenic reactions are then performed using an automated INTELLIPATH FLX system (Biocare Medical, Pacheco, CA, USA). Once the staining is complete, specimens are counterstained with Hematoxylin (Fisher Scientific, Hampton, NH, USA) and mounted using a xylene based mounting medium (Lecia Biosystems, Buffalo Grove, IL, USA).

### Flow Cytometry

Single-cell suspensions were generated from whole prostate tissues combining all prostate lobes using a modified procedure previously described (36). Briefly, whole prostates (combining all lobes) were dissected under sterile conditions from euthanized mice and collected in 1X HBSS buffer containing 5mM EDTA (Life Technologies Corporation, Grand Island, NY, USA) and 2% FBS (Hyclone, South Logan, UT, USA). They are incubated in a shaker for 15 min. at 37°C to loosen the tissue. Then the tissues were spun down at 100xg and minced with fine scissors. The tissues are then dissociated by shaking for 45 min. at 37°C in a 0.4 μm filtered solution of 0.5 mg/mL collagenase D (Roche, Indianapolis, IN, USA), 1 Unit/mL Dispase (Stemcell Technologies, Vancouver, BC, Canada), and 0.1 mg/mL DNase I (Sigma, St. Louis, MO, USA) in 1X HBSS. Digestions were subsequently filtered through a 40 μm nylon mesh and washed with 1X PBS twice before counting and proceeding to stain the cells. After washing, cells were incubated with Zombie UV^TM^ fixable viability kit (BioLegend®, San Diego, CA, USA). Following which, cells were washed in FACS buffer (2% FCS in PBS) and staining performed using the following conjugated antibodies: Brilliant Violet^TM^ (BV)510CD11c (catalog # 117353, RRID:AB_2686978), BV570-CD8 (catalog # 100740, RRID:AB_2563055), BV650-CD3 (catalog # 100229, RRID:AB_11204249), AlexaFluor700-CD45 (catalog # 103128, RRID:AB_493715), PerCP-B220 (catalog # 103234, RRID:AB_893353), PE/Dazzle^TM^594-CD4 (catalog # 100566, RRID:AB_2563685) (BioLegend®, San Diego, CA), and APC/Cy7-CD11b (catalog # 557657, RRID:AB_396772; BD Biosciences, San Jose, CA, USA). Samples were run on a BD LSRFortessa™ (BD Biosciences, San Jose, CA, USA) cytometer and analyzed using FlowJo™. The gating strategy for all the samples is shown in Fig. 7A.

### Real-time quantitative reverse-transcriptase PCR

Total RNA was isolated using TRIzol^TM^ reagent (Life Technologies Corporation, Grand Island, NY, USA), and cDNA synthesis, starting with 1 μg of total RNA, was performed with random hexamers using High-Capacity cDNA Reverse Transcription Kit (Life Technologies Corporation, Grand Island, NY, USA) per the manufacturer’s instructions. Primers for quantitative PCR (qPCR) were created for the RNA of interest using the NIH online primer blast tool. Reverse Transcription – Quantitative Polymerase Chain Reaction (RT-qPCR) reactions were performed using SsoAdvanced™ universal SYBR^®^ green (Bio-Rad, Hercules, CA, USA) and run on the CFX Connect (Bio-Rad, Hercules, CA, USA) platform. A full table of primer sequences is included in Table 1. The data were analyzed by the ^CT^ method (37), and normalized to GAPDH as the housekeeping gene. The data are represented as fold change normalized to the average expression of the gene of interest in their respective control group.

**Table 1:**
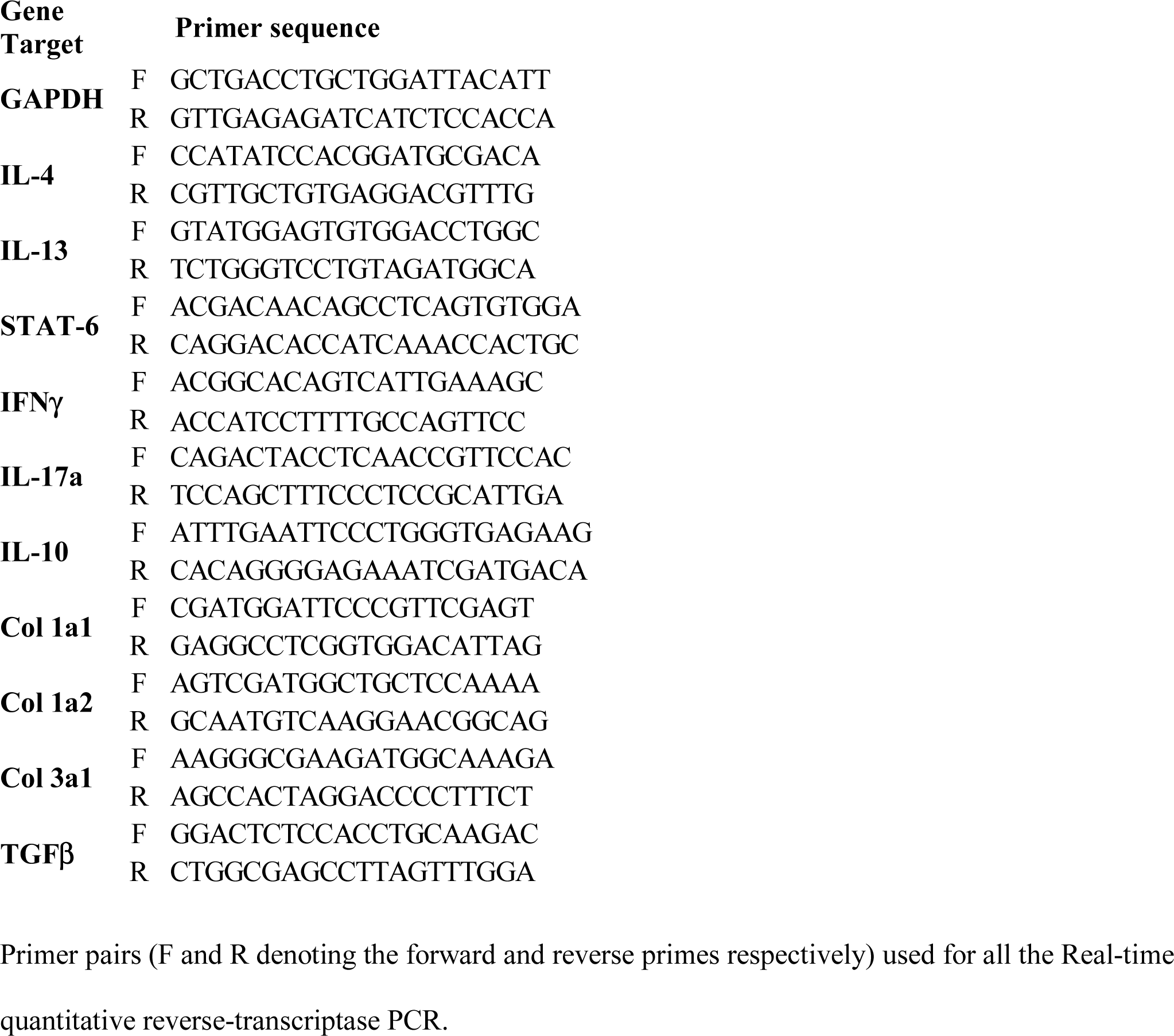
List of primer pairs used for quantitative real-time PCR.

### Western blotting

Frozen prostate samples were lysed in 1X RIPA lysis buffer (Santa Cruz Biotechnology, Dallas, TX, USA) containing completeLJEDTA protease inhibitor cocktail and phosSTOP phosphatase inhibitor (Millipore-Sigma, Burlington, MA, USA). Cell lysates were cleared by centrifugation at 14,000 rpm for 30 min at 4°C, and insoluble debris was discarded. Proteins were separated by SDS-PAGE on Criterion 4–20% mini gels (Bio-Rad, Hercules, CA, USA), transferred to polyvinylidene fluoride membranes (Bio-Rad, Hercules, CA, USA), blocked, and probed with the respective Abs. Immunoblotting was performed using the following antibodies - mouse anti-phospho myosin light chain 2 (Ser19) (catalog # 3675, RRID:AB_2250969), rabbit anti-human myosin light chain 2 (D18E2) [with cross-reactivity to mouse] (catalog # 8505, RRID:AB_2728760; Cell Signaling Technology, Danvers, MA, USA), and goat anti-human GAPDH [with cross-reactivity to mouse] (catalog # AF5718, RRID:AB_2278695; R&D Systems, Inc., Minneapolis, MN, USA); and developed using SuperSignal^TM^ west -pico or -femto chemiluminescence kit (Thermo Fisher, Hampton, NH, USA). The protein bands were quantified using the National Institutes of Health ImageJ software package and expressed as values of phosphorylated proteins normalized to total species and then to GAPDH levels.

### Statistical analyses

Statistical analyses were performed using GraphPad Prism™ (GraphPad Software, San Diego, CA, USA). Statistical tests utilized in each experiment, technical replicates, biological replicates, independent repeat experiments performed, and murine *n* values are indicated in figure legends (in most figures, each dot represents individual animals). Data are represented as the mean ± standard deviation (SD) or mean ± standard error of the mean (S.E.M.) as appropriate. # p < 0.1, * p < 0.05, ** p < 0.01, *** p <0.001, **** p < 0.0001.

## RESULTS

### Elevated mast cell numbers are observed in human BPH tissues

Prostate inflammation and fibrosis are linked to the development and progression of BPH (7–9, 11). Previous studies in the field of BPH have also shown that mast cells might be involved in the disease in human patients (7, 9, 22, 38, 39). To validate this information in our setting; fibrosis, as observed by collagen deposition captured by picrosirius staining, was examined in prostate tissue biopsy sections from human BPH patients and controls obtained from US Biomax, Inc. (Cat# PR632). Patients with BPH show an increase in immune cell infiltration as compared to controls (as shown in the company product specification images). Extracellular collagen deposition, a marker for fibrosis in tissue(40), showed an increase but non-significant trend (p value <0.1) in the prostate tissue of BPH patients compared to normal controls (Fig. 1, A&C**; C –** representative picrosirius images comparing surgical BPH and control normal prostate tissue). Mast cell numbers were quantified by immunostaining for mast cell tryptase in prostate tissue biopsy sections from BPH patients and controls. Our results show a significant increase in mast cells in prostate tissues from patients with BPH compared to controls (Fig. 1, B&D**; D –** representative IHC images comparing surgical BPH and control normal prostate tissue). This observation implies that mast cells may affect BPH development and progression.

**Fig. 1:**
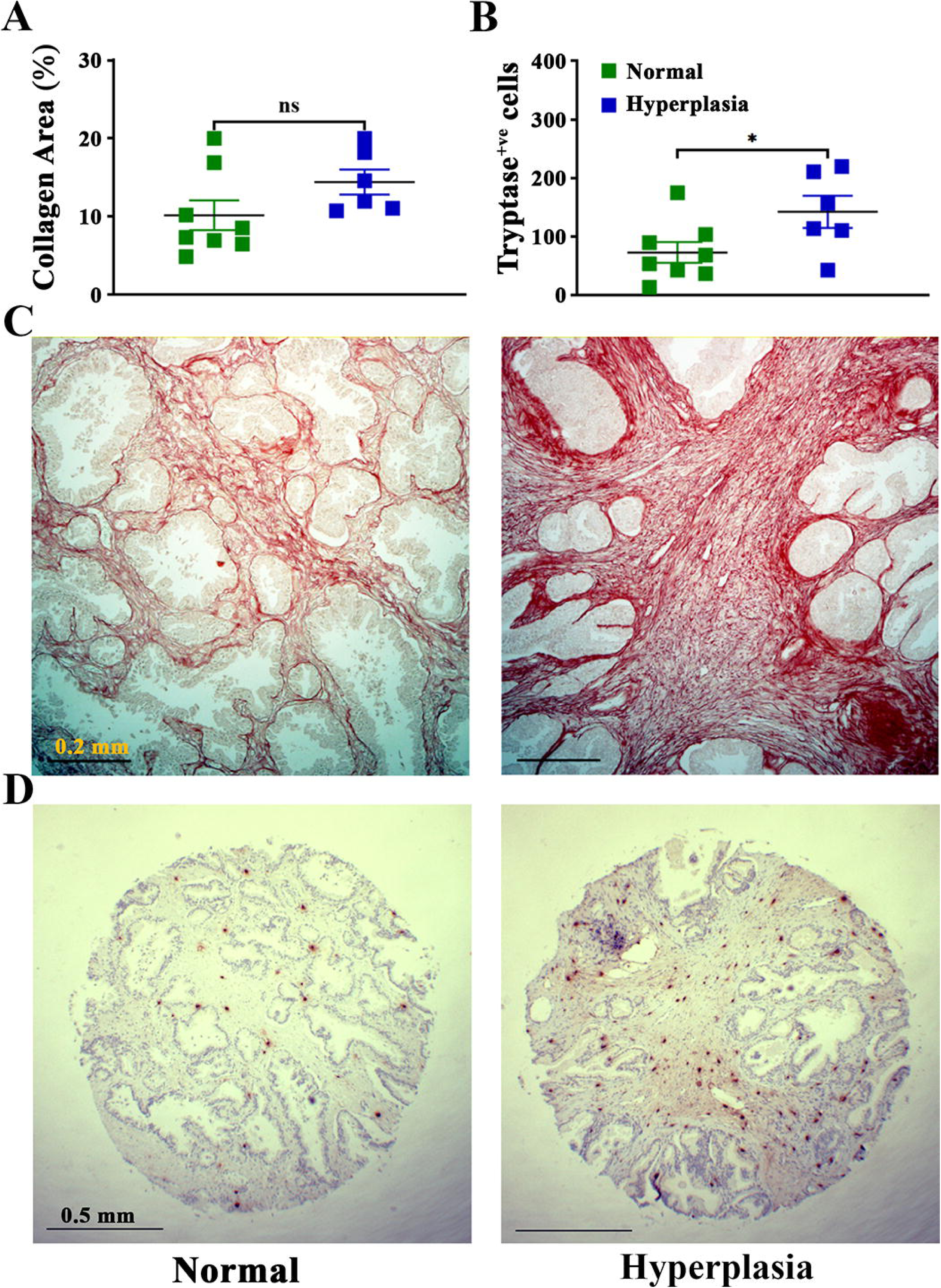
Increased fibrosis and mast cells numbers in prostates of BPH patients compared to normal control patients. Picrosirius staining for extracellular collagen deposition in human tissues **(A & C)** and IHC for human mast cell tryptase was used to assess numbers of mast cells **(B & D)** (from US Biomax, Inc.; catalog # PR632). **(A)** Quantification of percentage of extracellular collagen deposition in hyperplasia - BPH and normal prostate tissues using NIH Image J software s, and **(B)** quantitated data for number of human mast cell tryptase positive mast cells in hyperplasia - BPH and normal prostate tissue. Each data point represents an individual patient tissue data, which is derived by averaging the counts and collagen contents for 3 different tissue biopsies for each patient (Mean ± SEM; *, p < 0.05; unpaired t-test). **(C, D)**, representative images of prostate sections from normal control (left), and hyperplasia - BPH (right) patients under 10X magnification for picrosirius staining **(C)**, and 4X magnification for human mast cell tryptase **(D)**.

### Urinary dysfunction in the CP1-induced mouse model of LUTS is associated with activation of mast cells and an increase in mast cell numbers in the prostate

Our laboratory has previously demonstrated that intraurethral instillation of CP1 induces chronic prostate inflammation and fibrosis leading to the development of voiding dysfunction in mice (28, 30). Here, in addition to confirming that instillation of CP1 induces voiding dysfunction, we determined whether there were associated changes in mast cell numbers and activation status in the prostates of mice.

To determine effects on voiding behavior we utilized the urovoid system, a noninvasive approach to assess conscious urinary voiding behavior of mice (31). Representative data from the Urovoid analysis software that was utilized to determine the inter-micturition interval (IMI) and void mass (volume) per void in CP1-infected C57BL/6 animals over time (Fig. 2A). Following instillation with CP1, we observed a significant reduction in the average number of urinary voids at days 5, 14, and 35 compared to control non-infected mice. (Fig. 2B). Conversely, the average urinary void mass and IMI were significantly increased in mice at days 5, and 14 (and not at day 35) following instillation with CP1 compared to control mice (Fig. 2, C-D). The urine production rate (UPR) for both groups of mice remain the same over time, indicating that the mice do not have an infection induced intrinsic deficit in the production of urine (Fig. 2E).

**Fig. 2:**
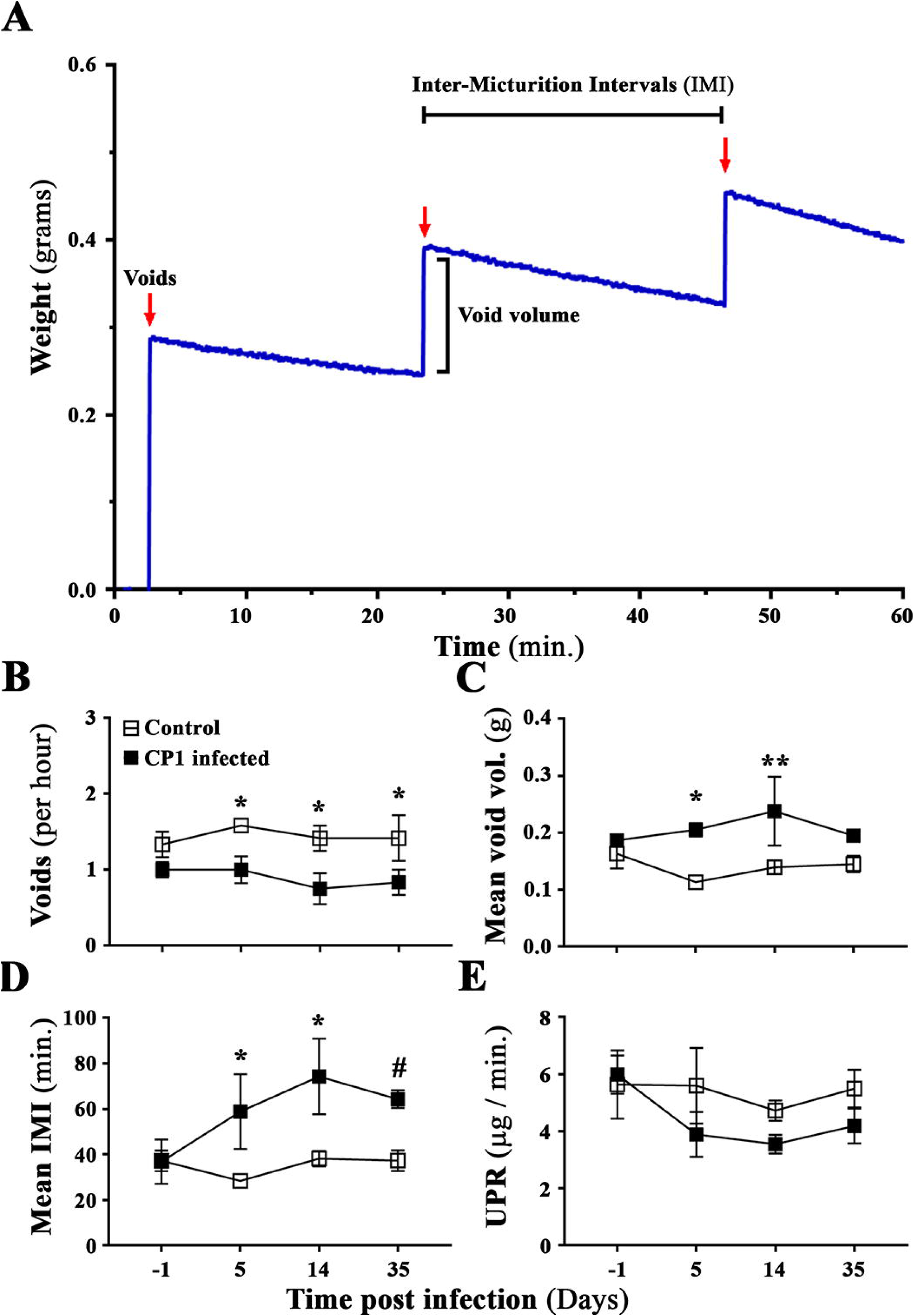
An *E. coli* (CP1)-induced mouse model of LUTS recapitulates and induces urinary dysfunction in mice. **(A)** a representative sample data graph from the Urovoid Analysis software showing the voiding measurements collected using the Urovoid system and how each of the parameters are calculated. The representative data shows the urine volume collected in weight (grams) over time (60 min.). The red arrows mark each individual void, the increase in weight per void is marked as void volume, and the time between each void is marked as Inter-micturition interval (IMI). **(B)** average number of voids per hour over a period of 4 hours; **(C)** average volume of urine per void (calculated for each animal individually); **(D)** average amount of time, in minutes, between each micturition event; and **(E)** average urine production rate (UPR) as defined as the volume of urine collected over the 4-hour period of time (indicating changes in the ability to produce urine) in control and CP1 infected C57BL/6 a day before and at days -5, -14, and -35 post-infection. (Mean ± SEM; N = 4 mice per group; ^#^ p < 0.1, * p < 0.05, ** p < 0.01; two-way ANOVA Fisher’s LSD test)

To assess the role of mast cells, we next examined the prostates of CP1-infected mice to determine the numbers and activation status of mast cells. Prostate sections from CP1 infected mice and control mice were subjected to toluidine blue staining to assess mast cell numbers and activation status in the lobes of the prostate. We observed a significant increase in the mast cell numbers in the prostate sections at days 5, 14, and 35 compared to control mice, and in particular, the increase in mast cell numbers was most striking in the dorsolateral lobe of the prostate (Fig. 3A). Furthermore, we observed an accumulation of mast cells in the prostates over the 35-day time period post CP1 instillation. We also assessed the activation status of these mast cells by quantitating the number of degranulated mast cells as compared to resting mast cells and determined the percentage of activated mast cells in the prostate sections. Following CP1 instillation, we observe a significant increase in the percentage of activated mast cells in the prostate sections as days 5, 14, and 35 compared to control mice (Fig. 3B) suggesting that CP1 infection causes increased degranulation of mast cells. Fig. 3C shows representative toluidine blue staining images of dorsolateral prostate tissues from control and CP1 infected C57BL/6 at day 35 post-infection showing increased mast cell infiltrates as well increased activated mast cells in the prostate tissues of CP1 infected mice.

**Fig. 3:**
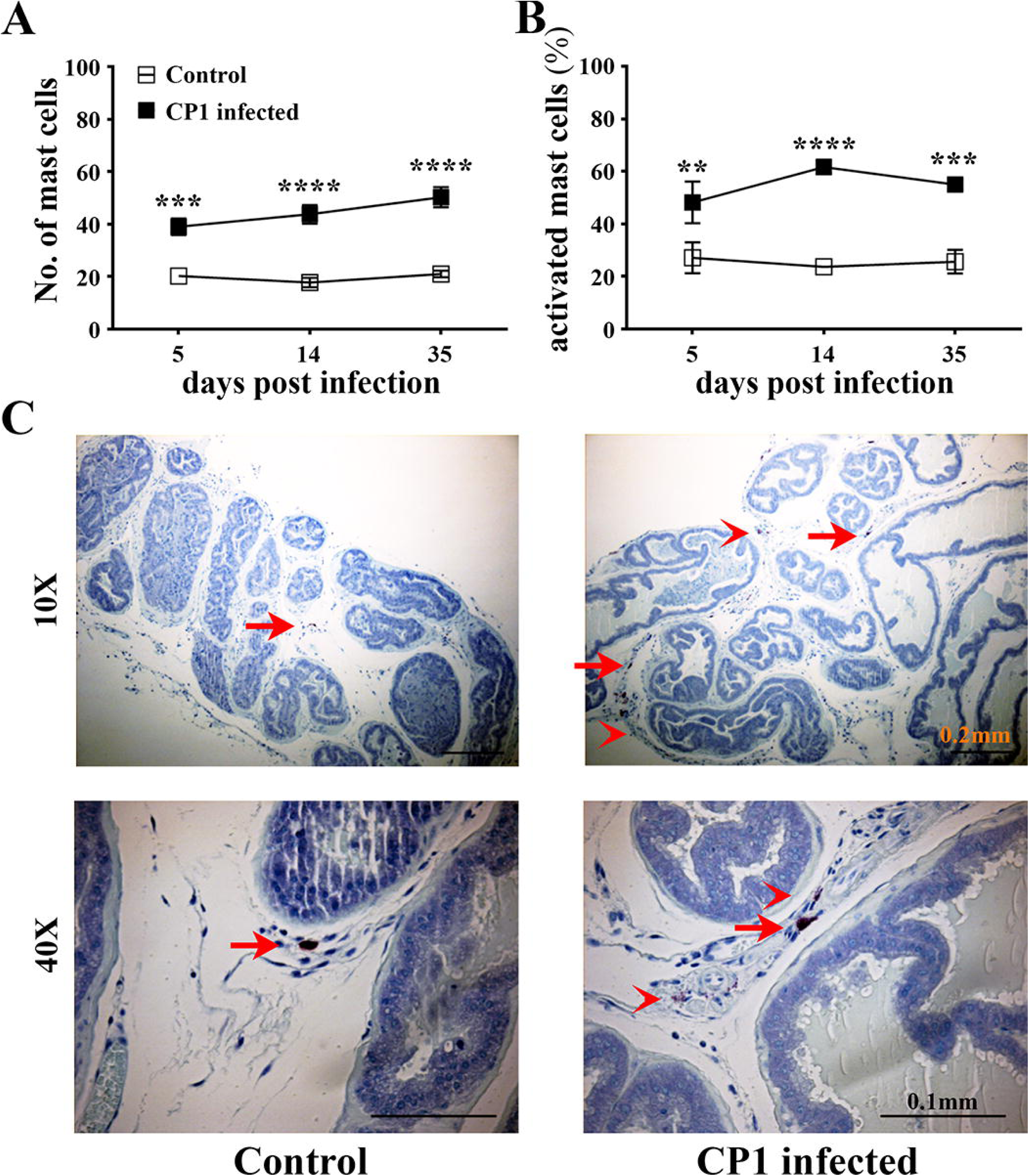
CP1 infection triggers an increase mast cell numbers and its activation in the prostate of mice. **(A)** Total numbers of mast cells and **(B)** percentage of activated (fully and partial combined) mast cells in toluidine blue stained sections of mouse prostate lobes harvested at 5-, 14-, and 35-days’ post CP1 infection (Mean ± SEM; N = 4 mice per group; ** p < 0.01, *** p < 0.001, **** p < 0.0001; two-way ANOVA Fisher’s LSD test). The mast cell numbers and percentage of activated mast cells were determined by counting and averaging three sections of the mouse prostate stained with toluidine blue for each sample. **(C)** representative images of toluidine blue stained dorso-lateral prostate sections from control PBS instilled (left) and CP1 infected (right) mice at 35-day post instillation imaged at 10X and 40X magnification. Note the increased degranulation of mast cells (as denoted by less intracytoplasmic granular staining of mast cells) is marked by red arrow heads, while resting mast cells (with intact cytoplasm and membrane) is marked by red arrows.

### Mast cell stabilizer and histamine 1 receptor inhibition prevents mast cell activation and alleviates urinary dysfunction in CP1 infected mice

To assess the role played by activated mast cells in the development and progression of BPH, we hypothesized that preventing mast cell activation and mast cell downstream signaling histamine signaling, as previously described, might be an effective strategy at reducing or reversing markers of pathology (33). To examine this, mice were administered a combination of a mast cell stabilizer (MCS), cromolyn sodium salt (CrS), and a histamine receptor 1 antagonist (H1RA), cetirizine di-hydrochloride (CeHCl) intra-peritoneally for 10 days daily starting at either 5- or 25-days (early and late respectively) post-infection (Fig. 4A).

**Fig. 4:**
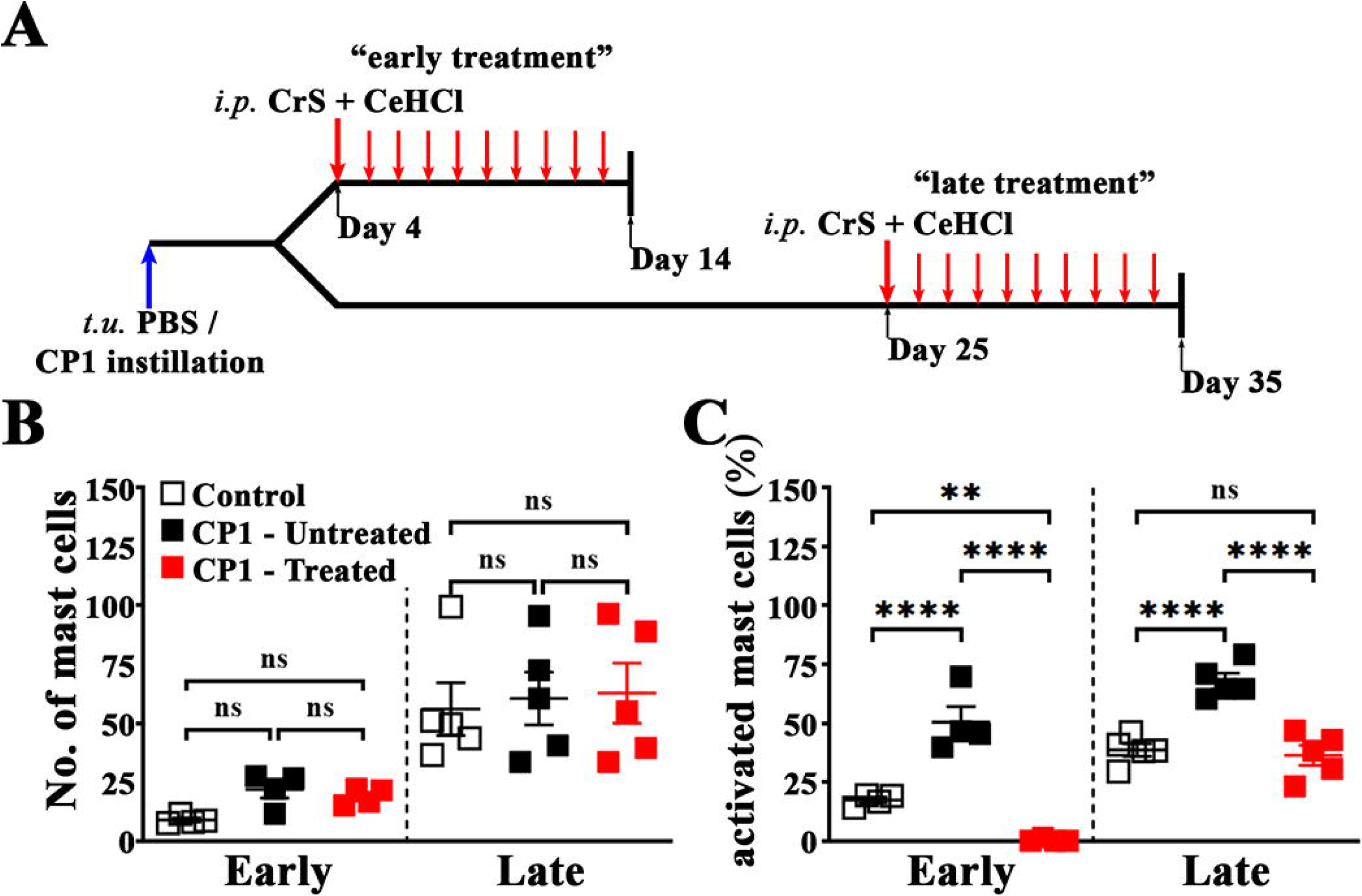
Combination treatment of cromolyn sodium and cetirizine di-hydrochloride inhibits mast cell activation in the prostates of CP1 infected mice. **(A)** Schematic representation of treatment of mice with cromolyn sodium (CrS) and cetirizine di-hydrochloride (CeHCl). Mice given transurethral (*t.u.*) instillations of PBS or CP1 were treated intra-peritoneally (*i.p.*) with the combination of CrS + CeHCl daily for 10 days from day 4 or day 25 (“early treatment” and “late treatment” respectively) before assessment of the effects of combination treatment on various parameters. **(B)** Total numbers of mast cells and **(C)** percentage of activated (fully and partial combined) mast cells in toluidine blue stained sections of mouse prostate dorso-lateral lobes harvested at 14 (early) and 35 (late) days’ post CP1 infection (Mean ± SEM; each dot represents an individual mouse; ** p < 0.01, *** p < 0.001, **** p < 0.0001; one-way ANOVA Fisher’s LSD test). The mast cell numbers and percentage of activated mast cells were determined by counting and averaging three sections of the mouse prostate lobes stained with toluidine blue for each sample.

To validate that this combination therapy is effective in preventing mast cell activation, we assessed the numbers and activation status of mast cells in prostate lobes from CrS + CeHCl treated CP-infected mice as compared to CP1-infected mice using toluidine blue staining of prostate sections. We observed that in both the “early treatment” and the “late treatment” groups, while there are little changes in the mast cell numbers in the combined prostate lobes of CrS + CeHCl treated CP1 infected mice as compared to CP1 infected mice, we observe a significant decrease in the % of activated mast cells following both “early” and “late” treatment (Fig. 4, B-C).

We next assessed the voiding behavior in CP1-infected mice post mast cell and histamine 1 receptor inhibition using the urovoid system. Similar to earlier observations (Fig. 2), following CP1 instillation mice experienced urinary dysfunction as observed by decreased average number of urinary voids in both “early” and “late” treatment groups (Fig. 5A), along with increased mean void mass and IMI (Fig. 5, B-C). Following 10-day i.p. administration of the combination of CrS + CeHCl, we observed the average number of urinary voids in CP1-infected mice were significantly increased comparable to that of control mice in the “late” treatment group (Fig. 5A). Furthermore, in both the “early” and “late” treatment groups, we observed a significant decrease in IMI that was similar to control mice (Fig. 5C).

**Fig. 5:**
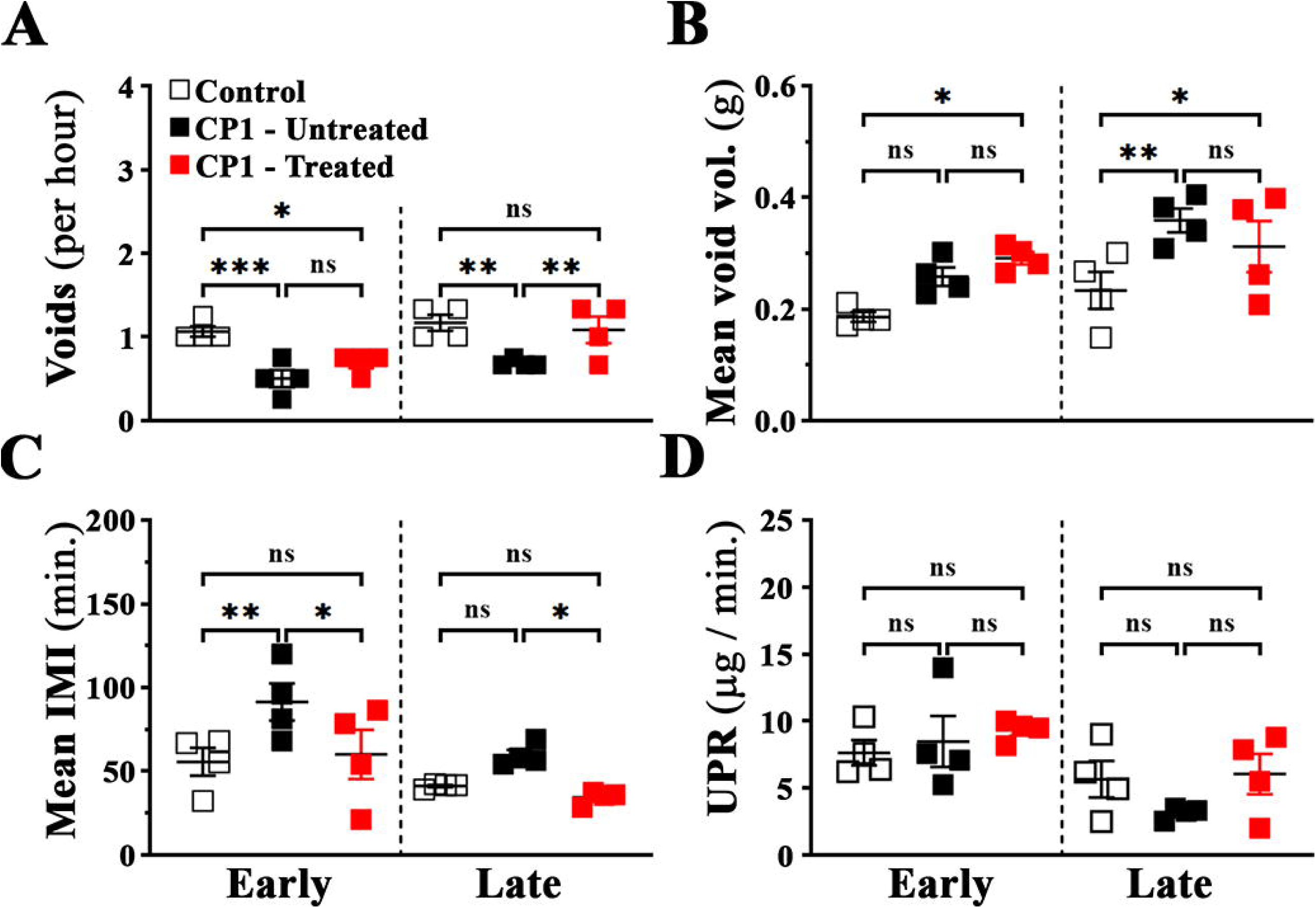
Mast cell inhibition alleviates urinary dysfunction in CP1 infected mice. **(A)** average number of voids per hour over a period of 4 hours; **(B)** average volume of urine per void (calculated for each animal individually); **(C)** IMI as measured by average amount of time, in minutes, between each micturition event; and **(D)** average urine production rate (UPR) in control, CP1 infected mice, and CrS + CeHCl treated CP1 infected mice at 14 (“early”) and 35 (“late”) days’ post CP1 infection (Mean ± SEM; each dot represents an individual mice; ** p < 0.01, *** p < 0.001; one-way ANOVA Fisher’s LSD test).

Interestingly, the average urinary void mass of CP1-infected mice treated with CrS + CeHCl did not show any difference as compared to untreated CP1-infected mice in both “early” and “late” treatment groups (Fig. 5B). The UPR for all the groups of mice remains the same at “early” and “late” time points (Fig 5. E). These data suggest treatment for inhibition of mast cell degranulation along with histamine 1 receptor inhibition (here in referred to as mast cell inhibition) leads to significant alleviation in CP1-induced urinary dysfunction in mice.

### Mast cell inhibition ameliorates fibrosis in prostates of CP1 infected mice

To understand the mechanism by which mast cell inhibition alleviates urinary dysfunction, we assessed inflammation and fibrosis in MCS + H1RA treated CP1-infected mice. We assessed inflammation in prostate tissue from CrS + CeHCl treated CP1-infected mice, CP1 infected mice, and control mice by staining sections from the prostate lobes using H&E. As previously shown (28), we observed that CP1 instillation in C57BL/6 mice triggers a modest but significant level of inflammation in the prostate sections from the dorso-lateral lobe of mice at day 35 (“late” group) compared to control mice (Fig. 6A). Upon mast cell inhibition, we observed no significant changes in the inflammation scores in both the “early” and “late” treatment group (Fig. 6A). Fig. 6C shows representative H&E images of sections from the dorsolateral lobe of prostate tissues from control, CP1-infected, and CP1-infected CrS + CeHCl treated C57BL/6 at day 35 post-infection showing infiltration of immune cells in the stroma of the prostates.

**Fig. 6:**
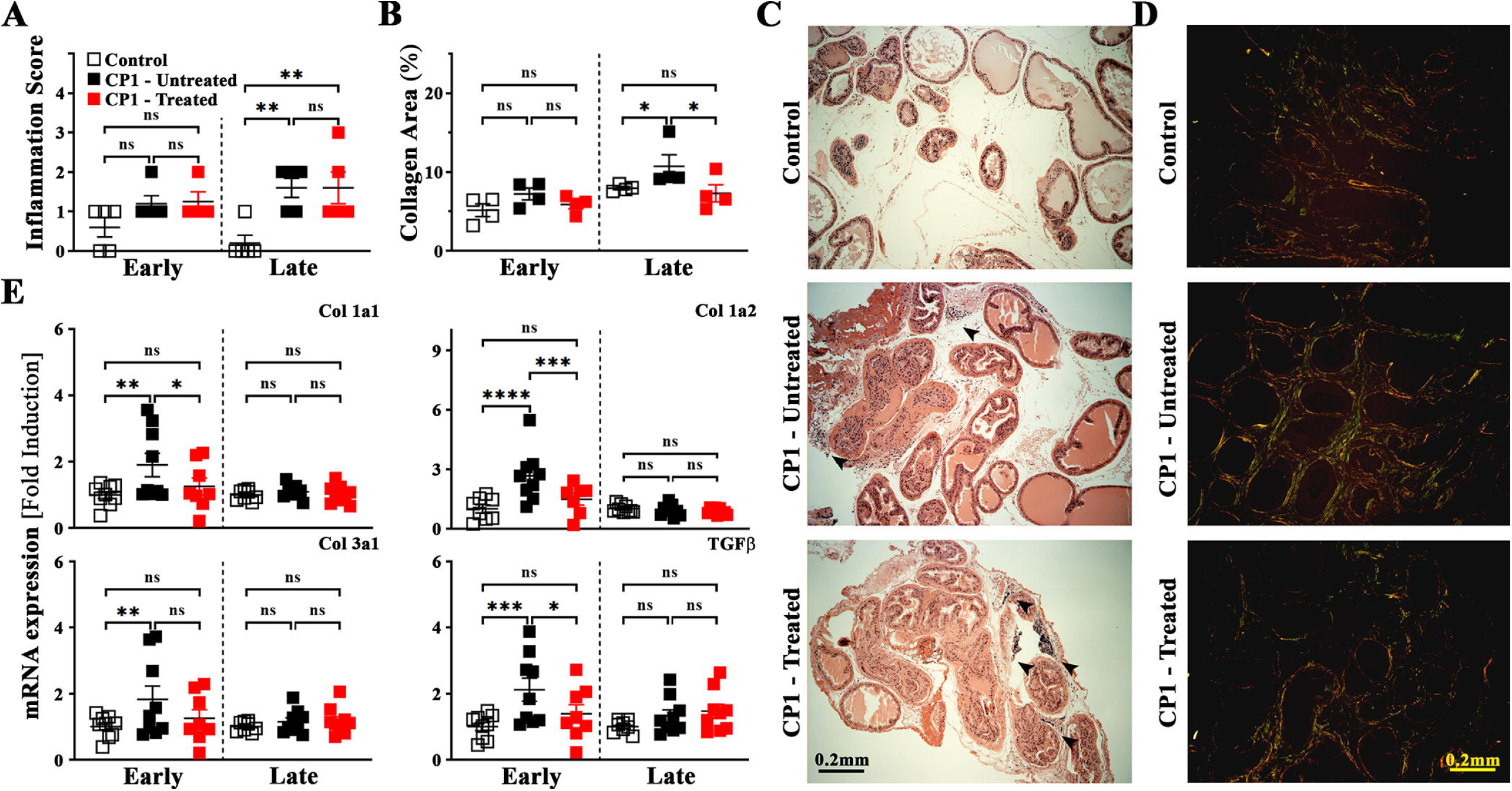
Inflammation and fibrosis is driven by mast cell activity in CP1-induced mouse model of LUTS. Quantitation of **(A)** inflammation score (as described in the materials and methods) using H&E staining, and **(B)** percentage of extracellular collagen deposition as assessed by picrosirius staining from sections of the dorso-lateral lobe of the prostates from control, CP1 infected mice, and CrS + CeHCl treated CP1 infected mice at 14 (“early”) and 35 (“late”) days’ post CP1 infection (Mean ± SEM; each dot represents an individual mouse; * p < 0.05, ** p < 0.01; one-way ANOVA Fisher’s LSD test). The inflammation score and percentage of collagen area were obtained by examining three independent sections of the mouse prostate lobes stained with H&E and picrosirius red staining respectively. Quantification of picrosirius staining was performed using NIH Image J software. Representative H&E images **(C)** and picrosirius images **(D)** of dorso-lateral prostate sections from control PBS instilled (top), CP1 infected (middle), and mast cell inhibitor treated CP1 infected (bottom) mice at 35-days post instillation imaged 4X magnification. The increased infiltration and immune cell foci is marked by black arrow heads **(C)**. **(E)** mRNA expression levels for pro-fibrotic genes Col -1a1, 1a2, and 3a1 as well as TGFβ were examined from RNA isolated from total prostates of mice from control, CP1 infected mice, and CrS + CeHCl treated CP1 infected mice at 14 (“early”) and 35 (“late”) days’ post CP1 infection. RNA was isolated, converted to cDNA, and subjected to real-time PCR analysis with the respective gene primers. The mRNA expression levels are normalized to GAPDH mRNA levels for each sample and the data are represented as fold change over control (Mean ± SEM; each dot represents an individual mouse; * p < 0.05, ** p < 0.01, *** p < 0.001, **** p < 0.0001; one-way ANOVA Fisher’s LSD test).

Next, we examined the extent of extracellular collagen deposition in each lobe of the infected mouse prostates. Previously we have shown that fibrosis, as observed by staining with picrosirius red, was significantly upregulated in the dorso-lateral lobe (and to a lesser extent in the ventral lobe) of the prostates of CP1 infected mice (28). Here, we observed similar results for extracellular collagen deposition in the dorsolateral lobes of prostates of CP1 infected mice at day 35 post infection (Fig. 6B). Therapeutic administration of CrS + CeHCl in CP1 infected mice (significantly at “late treatment” and to a lesser extent “early treatment”) is able to attenuate CP1 induced collagen deposition (Fig. 6B). Fig. 6D shows representative picrosirius images of sections from the dorsolateral lobe of prostate tissues from control, CP1-infected, and CP1-infected CrS + CeHCl treated C57BL/6 at day 35 post-infection showing collagen deposition.

Additionally, we performed qPCR for fibrosis-associated genes and pro-fibrotic markers on RNA extracted from the mouse prostate tissues. We examined expression levels of collagen -1a1, -1a2, and - 3a1 which are extracellular markers of fibrosis as well as TGFβ, a well-known pro-fibrotic signaling molecule (41). Upon CP1 instillation, we observed a significant upregulation of mRNA expression for all four markers in the prostates of CP1-infected mice compared to control mice at day 14 post-infection, as previously reported (28) (Fig. 6E). Interestingly, when CP1-infected mice were treated with CrS + CeHCl, RNA extracted from the prostates of these mice show significantly decreased levels of expression of Collagen -1a1, -1a2 as well as TGFβ in the “early treatment” group (Fig. 6E), but not for Collagen -3a1. While we do observe some mild upregulation in the mRNA of these four pro-fibrotic markers in the prostates of the CP1 infected mice at day 35 post-infection, this is not significant; and neither are there any significant changes upon treatment with CrS + CeHCl.

### Mast cell inhibition alters the immune cell skewing and inhibits type-2 cytokine expression in the prostates of CP1 infected mice

Activation of mast cells is known to release a plethora of mediators that are important in triggering the infiltration of immune cells as well as inducing tissue repair after injury (14, 15, 18, 20). Chemokines secreted from the mast cells acting within the prostatic epithelium and stroma provide signals that contribute to increased numbers of B and T lymphocytes and macrophages in the prostates of patients with BPH (42). To assess the effect of the mast cell combination therapy on immune cell infiltrates, flow cytometry was performed on prostates of CP1-infected mice, and immune cell populations were identified and gated as shown in Fig. 7A. As seen in Fig. 7B, at day 35 post CP1 instillation, we observed an increase in the numbers of immune cell infiltrates in the prostates of CP1-infected mice (as seen by the total numbers of CD45^+^ cells) which is significantly decreased to levels comparable to that of control mice upon treatment with CrS + CeHCl. We observed that CP1-infection significantly increased the numbers of total CD3^+^ T cells, as well as CD8^+^ T cells in the prostates of mice. Mast cell inhibition (treatment with CrS + CeHCl) significantly decreased the numbers of total CD3^+^ T cells as well as CD8^+^ T cells to numbers similar to those in control mice. Furthermore, we also observed that upon CP1 instillation, there is a significant increase in numbers of CD11b^+^ macrophages and a modest increase in CD11c^+^ dendritic cells in prostates compared to control mice, and upon treatment with CrS + CeHCl the numbers of CD11b^+^ macrophages significantly decreased to numbers similar to those of control mice (Fig. 7B). These observations were observed in the “early treatment” group as well, albeit to a lesser extent considering that CP1 instillation does not trigger immune cell infiltration early during infection of C57BL/6 mice (Table. 2). As seen in Table. 2, in the “early treatment” group upon CrS + CeHCl treatment of CP1-infected mice, we observe a significant decrease in the numbers of CD11b^+^ macrophages, as well as CD4^+^ and CD8^+^ T cells compared to CP1-infected mice.

**Fig. 7:**
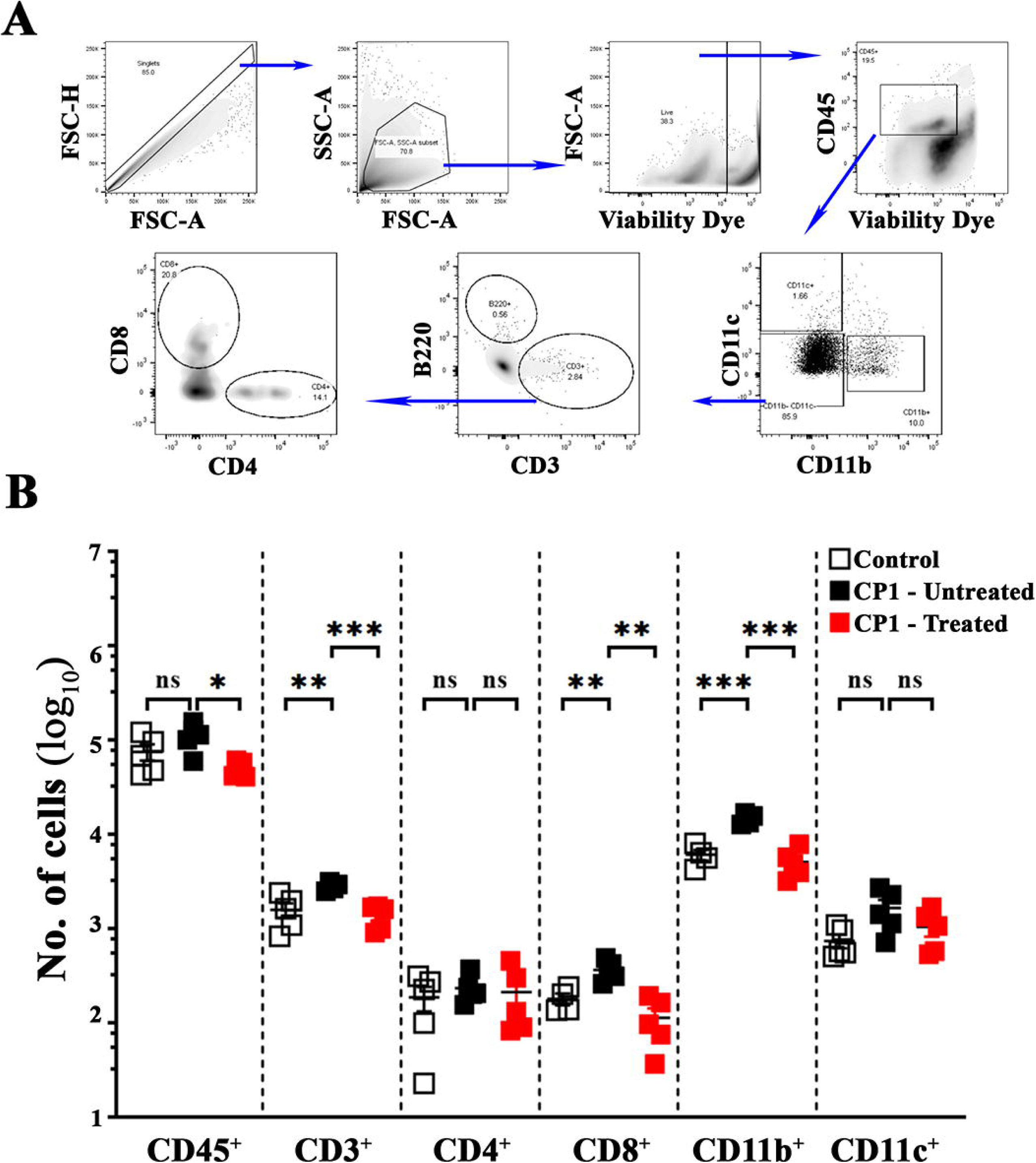
Mast cell inhibition mollifies alterations in CP1-induced immune cell infiltrates in the prostate of mice. **(A)** Flow cytometric gating strategy for analyzing the immune cell infiltrates in cell suspension prepared from whole prostates of mice. **(B)** Prostates isolated from control, CP1 infected mice, and CrS + CeHCl treated CP1 infected mice at 35 (“late”) days’ post CP1 infection were digested and total lymphocytes (CD45^+^ cells), CD3^+^ T-lymphocytes (as well as CD4^+^ and CD8^+^ T cells), CD11b^+^ monocytes/macrophages, and CD11c^+^ dendritic cells were stained with the respective fluorescent antibodies (as indicated in materials and methods) and analyzed by flow cytometry. Data represents total numbers of each cell type obtained from whole prostates using the tissue digestion protocol described in materials and methods (Mean ± SEM; each dot represents an individual mouse; * p < 0.05, ** p < 0.01, *** p < 0.001; one-way ANOVA Fisher’s LSD test).

**Table 2:**
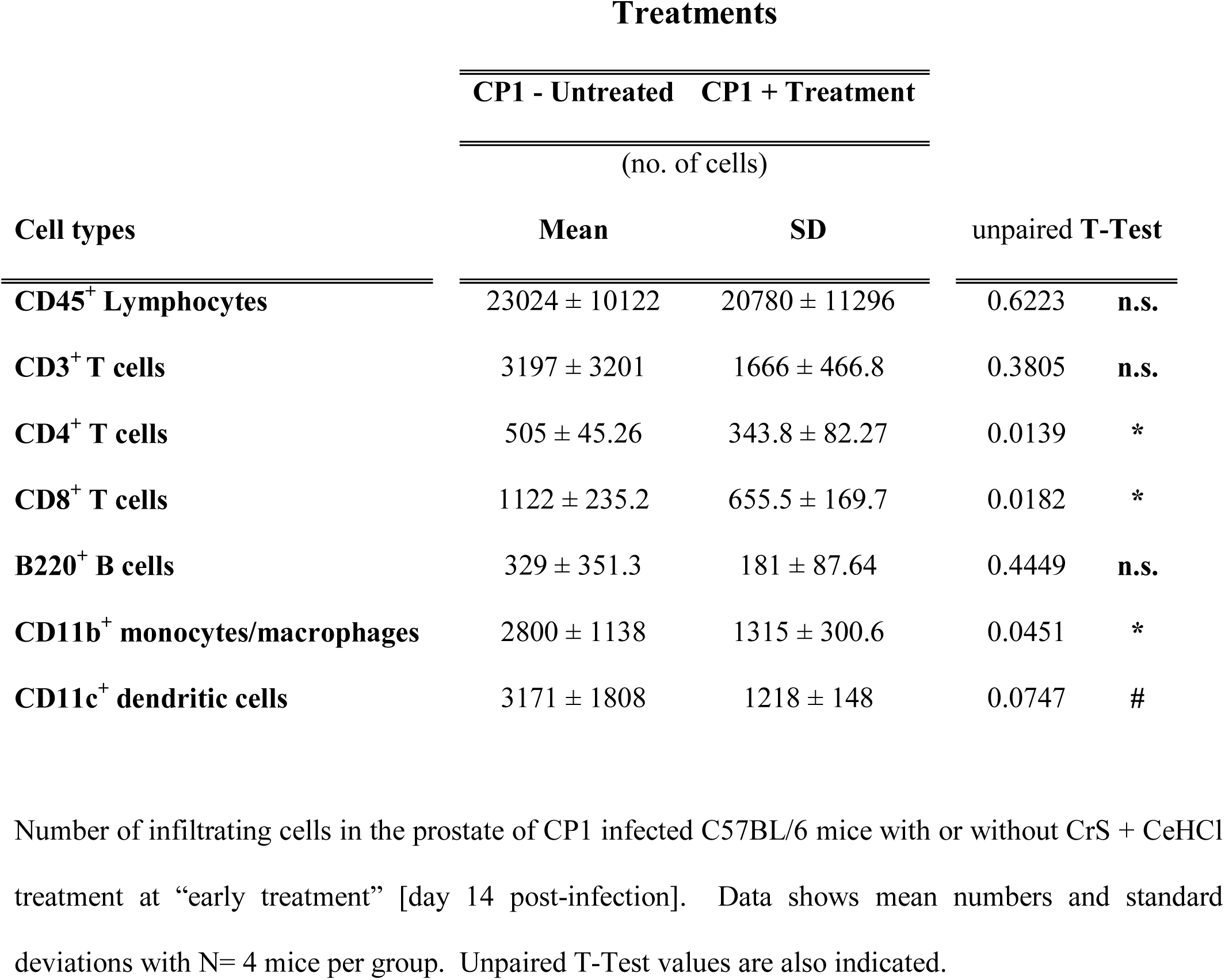
Immune cell infiltrates in the prostate of CP1 infected mice with and without CrS + CeHCl combination treatment.

| Cell types | Treatments |  | unpaired T-Test |  |
| --- | --- | --- | --- | --- |
|  | CP1 - Untreated | CP1 + Treatment |  |  |
|  | (no. of cells) |  |  |  |
|  | Mean | SD |  |  |
| CD45<sup>+</sup> Lymphocytes | 23024 ± 10122 | 20780 ± 11296 | 0.6223 | n.s. |
| CD3<sup>+</sup> T cells | 3197 ± 3201 | 1666 ± 466.8 | 0.3805 | n.s. |
| CD4<sup>+</sup> T cells | 505 ± 45.26 | 343.8 ± 82.27 | 0.0139 | * |
| CD8<sup>+</sup> T cells | 1122 ± 235.2 | 655.5 ± 169.7 | 0.0182 | * |
| B220<sup>+</sup> B cells | 329 ± 351.3 | 181 ± 87.64 | 0.4449 | n.s. |
| CD11b<sup>+</sup> monocytes/macrophages | 2800 ± 1138 | 1315 ± 300.6 | 0.0451 | * |
| CD11c<sup>+</sup> dendritic cells | 3171 ± 1808 | 1218 ± 148 | 0.0747 | # |

Number of infiltrating cells in the prostate of CP1 infected C57BL/6 mice with or without CrS + CeHCl treatment at “early treatment” [day 14 post-infection]. Data shows mean numbers and standard deviations with N= 4 mice per group. Unpaired T-Test values are also indicated.

We had previously reported that type-2 cytokine signaling plays an important role in driving fibrosis in our CP1-induced model of BPH associated LUTS (28). Further, type-2 cytokines have been shown to drive fibrosis in several different diseased conditions (40, 43, 44). To assess the impact of mast cell inhibition therapy on type-2 associated cytokines, qPCR was performed for cytokine transcripts from prostates obtained from control, CP1-infected untreated and CrS + CeHCl treated mice(45). As seen in Fig. 8, in the “early” treatment; and to a lesser extent in the “late” treatment groups; we observed a significant increase in the transcript levels of IL-4 and IL-13 (type-2 associated cytokines) as well as an upregulation of STAT6 (a key signal transduction molecule associated with Th2 polarization (46, 47)) in prostates of CP1-infected mice compared to controls. Upon treatment with CrS + CeHCl, the three markers show significantly reduced transcript levels compared to CP1-infected mice alone in the “early” treatment group. While the gene expression of other cytokines like IFNγ and IL-17 (associated with type-1 and type-3 immune responses, respectively), and IL-10, a negative regulator of T-cell activation, are significantly upregulated in the prostates upon CP1 instillation in the “early” treatment group. The combination treatment with CrS + CeHCl does not cause any significant changes to their transcript levels in comparison with CP1-infected mice (Fig. 8). These data show that the mast cell combination therapy inhibits type-2 cytokine skewing of the immune response in the prostates of mice induced by CP1 instillation.

**Fig. 8:**
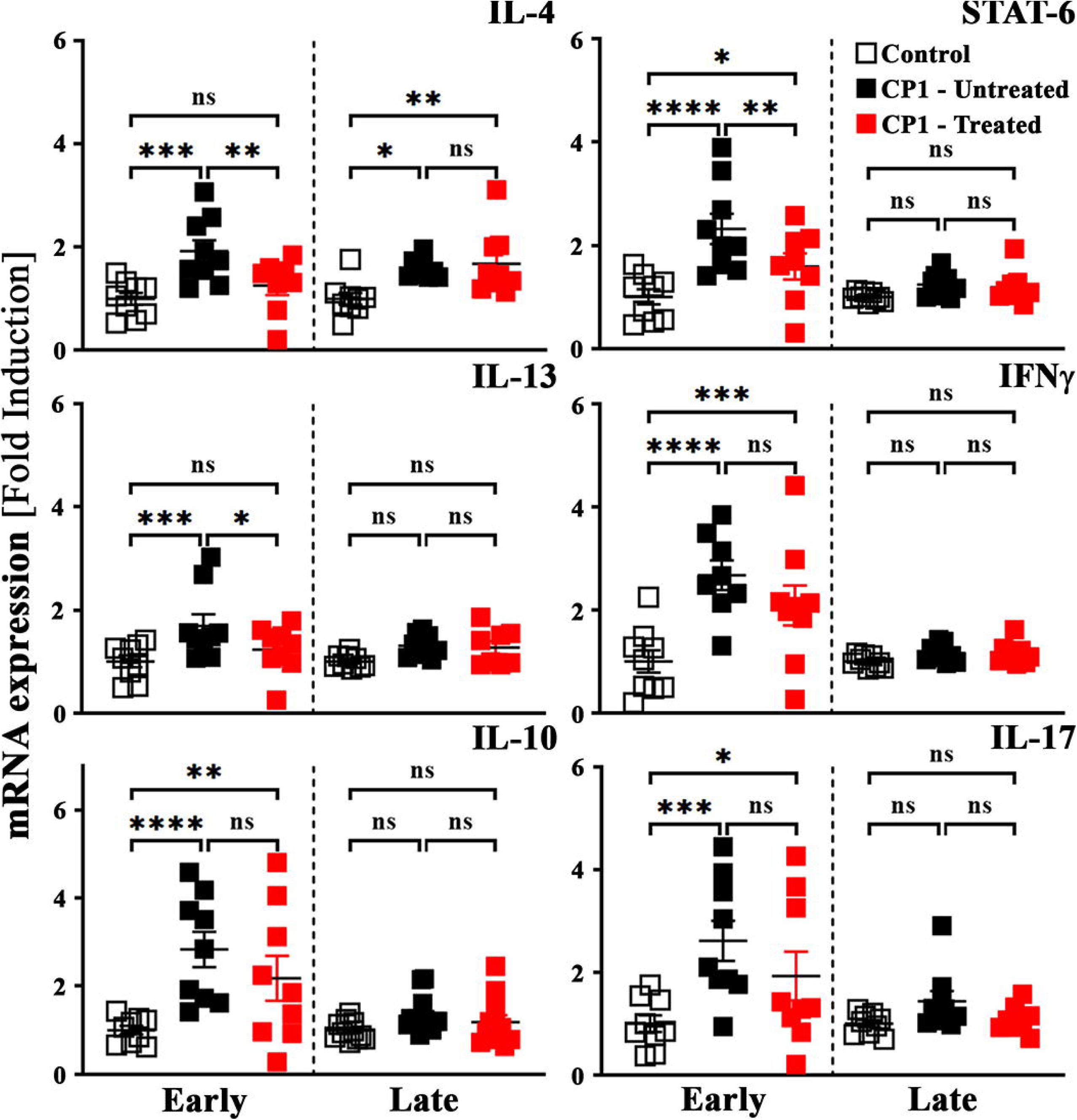
Prostates of CP1 infected mice treated with CrS + CeHCl show attenuated expression of type-2 cytokine gene expression. Whole prostates from control, CP1 infected mice, and CrS + CeHCl treated CP1 infected mice at 14 (“early”) and 35 (“late”) days’ post CP1 infection were lysed in TRIzol^TM^. RNA was isolated, converted to cDNA, and subjected to real-time PCR analysis with the respective gene primers. The mRNA expression levels are normalized to GAPDH mRNA levels for each sample and the data are represented as fold change over control (Mean ± SEM; each dot represents an individual mouse; * p < 0.05, ** p < 0.01, *** p < 0.001, **** p < 0.0001; one-way ANOVA Fisher’s LSD test).

### Mast cell inhibition attenuates prostate smooth muscle cell contraction in CP1-infected mice

Prostate stromal cell proliferation and smooth muscle contraction are crucial in the development and pathogenesis of BPH associated LUTS and have been a major target for treatment (48). Previously, we have shown that PAR2, a receptor for trypsin and the mast cell-derived serine protease and tryptase, plays an important role in the prostate smooth muscle cell contraction (27). As our combination treatment interferes with the release of mast cell proteases and tryptases, we next examined the consequences of the combination treatment on smooth muscle cell contraction. We assessed smooth muscle cell contraction by assessing the phosphorylation status of myosin light chain (MLC) -2, phosphorylation and de-phosphorylation of which regulates muscle contraction and relaxation respectively (49–51), in the prostates from control, CP1-infected untreated and CrS + CeHCl treated mice. As seen in Fig. 9A, upon CP1 instillation in mice, prostate lysates show an elevated level of MLC2 phosphorylation as compared to control mice in the “early treatment” group. Interestingly, in the “early treatment” group, we observed that mast cell inhibition causes a significant decrease in the levels of phosphorylated MLC2 in prostate lysates (Fig. 9A). Fig. 9B shows the quantitation by densitometry analysis of the western blot photomicrographs from multiple mice from both “early” and “late” treatment groups. In contrast to what was observed in the “early” treatment group, at later points post CP1 instillation, we observe no significant increase in the levels of phosphorylated MLC2 compared to control mice in prostate lysates. There are no significant differences in phosphorylated MLC2 levels in CP1 infected mice upon mast cell inhibition (Fig. 9B). The data suggest that the mast cell inhibition is effective in reducing smooth muscle cell contraction in the prostates of CP1 infected mice.

**Fig. 9:**
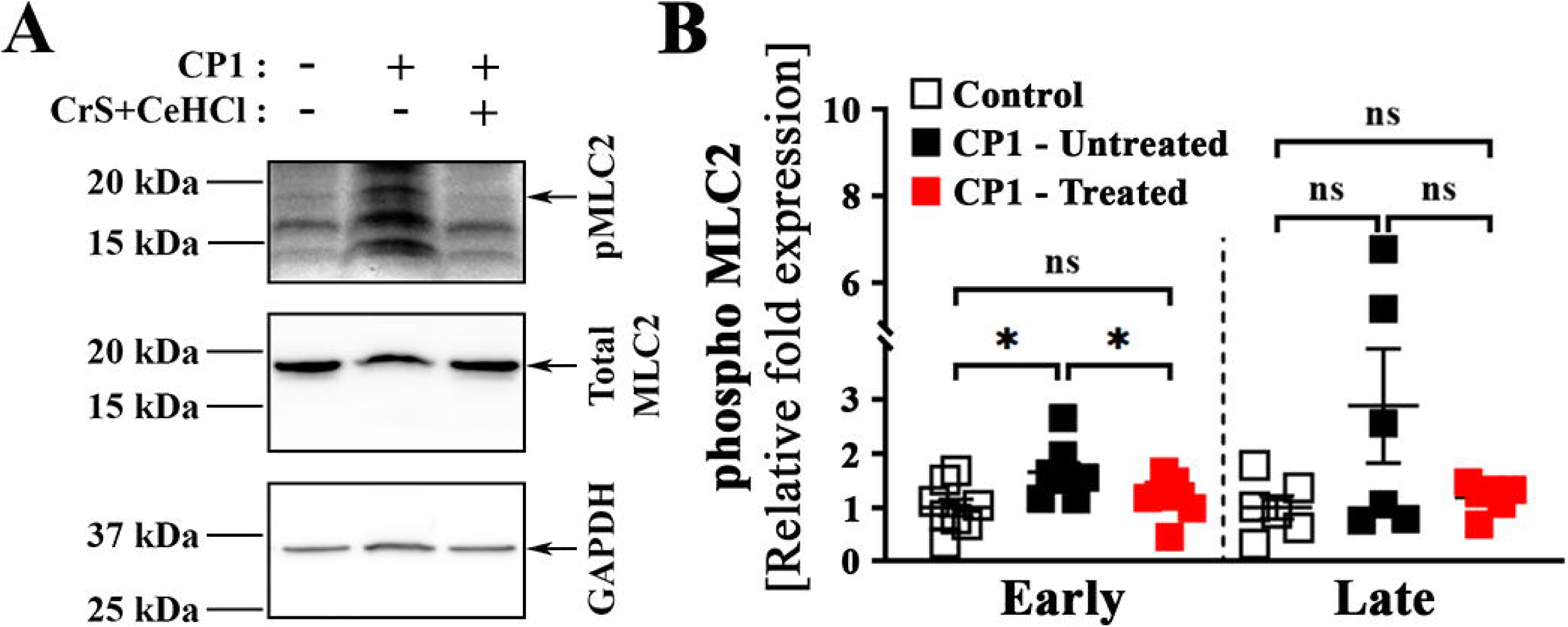
Mast cell inhibition during CP1-induced LUTS attenuates MLC2 phosphorylation in the prostates of mice. Whole prostates from control, CP1 infected mice, and CrS + CeHCl treated CP1 infected mice at 14 (“early”) and 35 (“late”) days’ post CP1 infection were lysed in 1X RIPA lysis buffer. **(A)** Whole-cell lysates were examined by Western blot analyses using the indicated antibodies, and a representative western blot for each group of mice at “early” times are shown. **(B)** quantifications of the Western blot data shown in **A** (n = 8 “early” and n = 7 “late”). The data plotted are arbitrary values obtained by normalizing the band intensity for the phosphorylated MLC2 to the band intensities of total MLC2 and then normalizing this to the band intensity of GAPDH, and the data are expressed as fold change over control (Mean ± SEM; each dot represents an individual mouse; * p < 0.05, ** p < 0.01, *** p < 0.001; one-way ANOVA Fisher’s LSD test).

## DISCUSSION

The pathogenesis of LUTS associated with BPH can be broken down into three facets: the epithelial compartment where hyperplasia of the epithelial cells occurs, the stromal compartment where immune cell infiltration and inflammation cause potential fibrosis, and lastly the smooth muscle compartment where smooth muscle cell contraction occurs (13, 38). An inflammatory insult, sterile inflammation, aging-related factors, or stress-induced hormonal changes, may trigger a dysregulation in any or all these compartments leading to the development and progression of LUTS associated with BPH (4, 5, 7–10). In this study, using a previously established *E. coli* (CP1) infection induced model of LUTS, we assessed the importance of mast cells in the development and progression of urinary dysfunctions.

Based on previous work, CP1 infection in C57BL/6 mice model triggers prostate inflammation, increased immune cell infiltration, urinary dysfunction, and fibrosis. Although the extent of inflammation and fibrosis, as well as the type of inflammation, are different in C57BL/6 mice as compared to NOD-ShiLJ mice, the CP1 infection in C57BL/6 mice does not induce pain (28, 29). The similarities between the urinary dysfunction associated with CP1 infection induced mouse model and LUTS in human BPH extends to increased presence of mast cells in the prostates (especially the dorso-lateral lobe) of the mice as well. This observation of the increased presence of mast cells in BPH tissue is not novel, but the importance of these mast cells has been relatively unappreciated (22, 52, 53).

Previously we demonstrated that CP1 infection in mice induced upregulation of type-1 (IFNγ), type-2 (IL-4, IL-5, IL-13) and type-3 (IL-17A) cytokines in the prostates and the draining lymph nodes (28, 30, 54). Furthermore, in Bell-Cohn *et al.*, we showed that this type-2 cytokine production from CD4^+^ T cells, via activation of the STAT6 pathway, was crucial in the development of fibrosis in the prostate of CP1 infected C57BL/6 mice. Here, our data further reinforces the idea that CP1 infection triggers type-1, type-2, and type-3 cytokine upregulation. More interestingly, we observed that combination treatment was specifically able to attenuate the production of type-2 (IL-4, IL-13) cytokines as well as STAT-6 gene expression. These findings can be explained by two different possible mechanisms. The combination treatment may directly affect the ability of the mast cells to produce and release type-2 cytokines upon activation and degranulation (14, 20). Secondarily, in synergy with the first proposed mechanism, the administration of H1RA acts as an immunomodulator in the Th1/Th2 imbalance in the diseased prostate and in this model dampens the production of type-2 cytokines by Th2 CD4^+^ T cells as well as preventsTh2 differentiation and infiltration (55, 56). The ability of the combination treatment to attenuate STAT-6 alongside IL-4 and IL-13 cytokine gene expression, suggests suppressing the downstream signaling of mast cell mediators through the administration of H1RA directly or indirectly skews the CD4^+ve^ T lymphocytes away from a Th2 cell type. This immunomodulatory effect can be seen in the dampening of fibrosis development in prostate tissue. In this current study, we were unable to evaluate the changes in the T helper cell subtypes because of the limitations in the numbers of cells extracted from the prostates of mice. However, studies are underway to assess the effects of this combination treatment in the modulation of the Th1/Th2 imbalance in the prostates and the draining lymph nodes.

An unexpected and very surprising observation from mast cell inhibition in the CP1-induced mouse model of LUTS is the ability of the combination treatment to inhibit increased presence of immune cells in the prostates of mice. A plethora chemokines and mitogens are released from mast cells in the context of inflammation, allergy, and infection (14, 18) and in some cases, chemokine release occurs independent of mast cell degranulation (57). The absence of increased immune cells in the prostates of CP1 infected mice upon mast cell inhibition could suggest that they either inhibit infiltration of immune cells in the prostates of these mice, or that they do not provide growth factors, cytokines and mitogens necessary for the proliferation of the immune infiltrates. In either scenario, our finding provides a crucial piece of evidence that mast cells might be playing an upstream role in this CP1 infection model of LUTS in mice which precedes immune cell skewing and fibrosis in the prostates of mice.

Mast cell released mediators including histamine, proteases, tryptases, chymases, and leukotrienes have been implicated to play a critical role in smooth muscle cell function, apoptosis, and contraction (15). These mast cell mediators act on PAR2 as well as histamine 1 receptors and play a role in triggering smooth muscle cell contraction (56, 58). Little is understood about the role of mast cell released factors in prostates of patients with BPH and its effects on smooth muscle cell contraction, however the relationship between mast cells, histamine 1 receptors, histamine 1 receptor antagonists, and its effect on airway smooth muscle cell contraction in regards to asthma has been well studied (59–62). PAR2 is a gene that is ubiquitously expressed in most tissues in the human body, and we had previously shown that PAR2 is expressed at high levels in the smooth muscle cells of both murine and human prostates. PAR2 activation by these mast cell proteases, triggers downstream Ca^2+^ dependent prostate smooth muscle cell contraction (27, 63). In the manuscript Paul *el al.*, we had discussed the possibility that a PAR2 inhibitors along with α-blockers (drugs that inhibit α-adrenergic receptor signaling) could be used in conjunction to treat urinary symptoms in patients with chronic prostatitis / chronic pelvic pain syndrome (CP/CPPS) and BPH. Our current observation provides another layer and possibly a more favorable therapeutic target upstream of PAR2 blockade, along with alleviating fibrosis and inflammation.

In summary, the key findings of our study show that mast cell numbers and its activity are increased in prostates of patients with BPH/LUTS and in CP1-infected mice showing symptoms of LUTS. We show that mast cell inhibition, through a combination of MCS and H1RA, alleviates the urinary dysfunction in CP1-infected mice. We observe that mast cell inhibition triggers decreased inflammation and fibrosis, attenuation in smooth muscle cell contraction, and a decrease in immune infiltrates in the prostates of CP1-infected mice (Fig. 10).

**Fig. 10:**
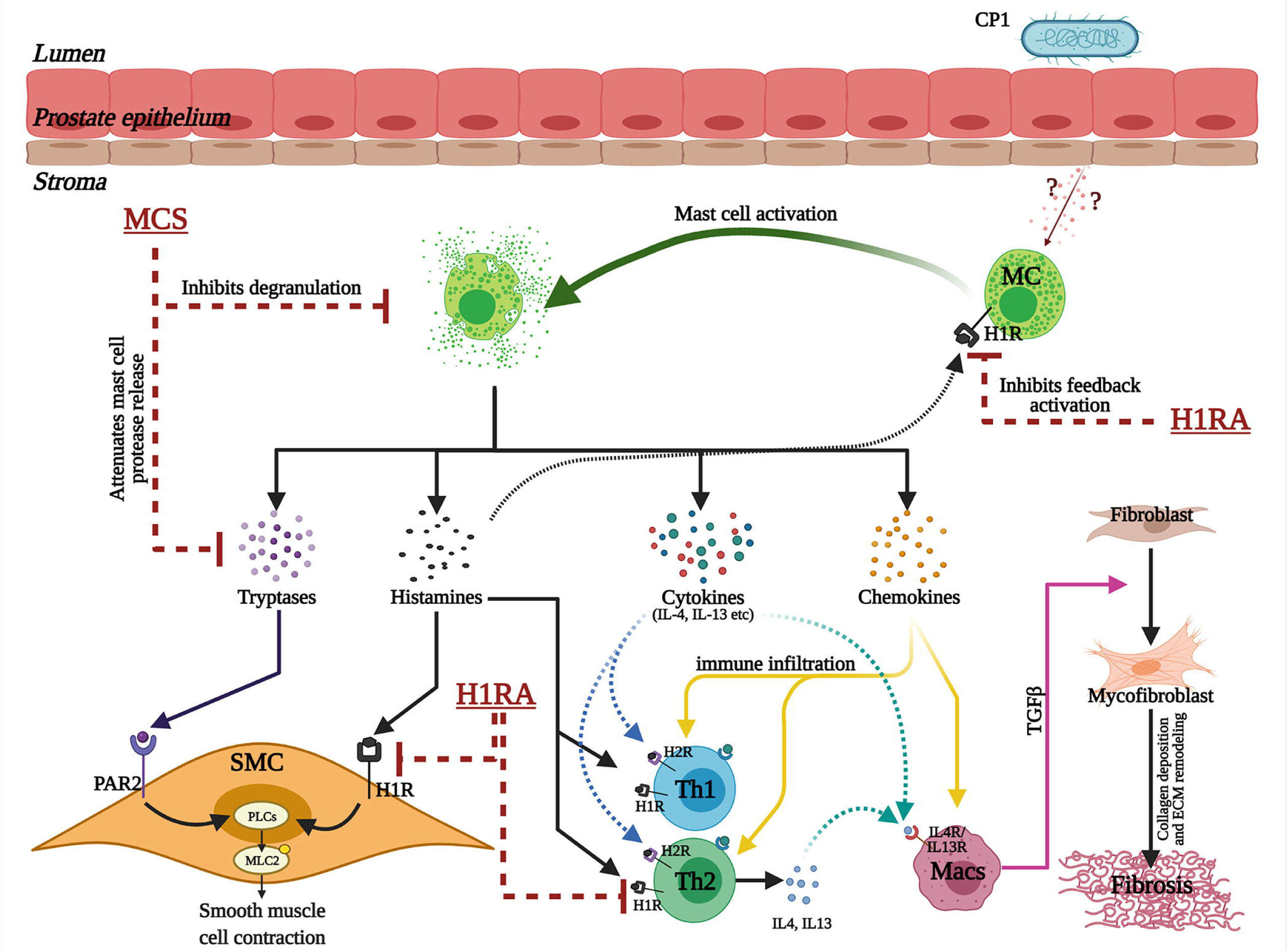
Schematic illustration of the crucial multifaceted role played by prostate mast cells in facilitating prostatic inflammation, fibrosis, and smooth muscle contraction in the development of lower urinary tract symptoms. Intraurethral instillation of CP1 in mice triggers epithelial tissue damage that induces the release of as yet unknown factors that lead to an increase in mast cell (MC) numbers and its activation in the mouse prostate stroma. The activated mast cells in turn release a plethora of factors which include tryptases (and other proteases), histamines, cytokines (including IL-4, IL-13), and chemokines. The tryptases and histamine released from mast cells act on smooth muscle cells (SMC), via protease-activated receptor 2 (PAR2) and histamine 1 receptors (H1R) respectively, leading to a PLC mediated regulation of smooth muscle cell contraction. Histamine released from activated mast cells also trigger a feedback loop to further activate mast cells through histamine receptors on mast cells. The chemokines released trigger increased immune cell infiltration and proliferation. The cytokines released from mast cells lead to activation of T helper type 1 (Th1), Th2, Th17 cells, dendritic cells and macrophages (Macs). The histamines released from mast cells in turn act on H1R and H2R on Th1 and Th2 cells modulating type-1/type-2 cytokine production leading to a hyper-inflammatory environment. IL-4 and IL-13 released from mast cells and Th2 cells act on macrophages (Macs) and cause macrophage polarization leading to release of TGFβ which causes primed resident fibroblasts to differentiate into myofibroblasts that release collagen and other matric proteins leading to extracellular matrix (ECM) remodeling and fibrosis. The combination treatment for mast cell inhibition which include mast cell stabilizer (MCS) and histamine 1 receptor antagonist (H1RA) act on multiple levels of these crucial aspects and regulate the tissue environment in a CP1-infected mouse prostate.

An in-depth assessment of the importance of mast cells in the development and progression of urinary dysfunction can be addressed using genetic knockout (mast cell-deficient W-sash c-kit mutant Kit^W-sh/W-sh^ mice) approaches (64–66). These approaches, however, could be confounded by the impact of mast cells on innate immune homeostasis. The combination therapy approach of administration of a MCS along with H1RA provides a model wherein we are able to dampen overactive mast cells in the prostate to alleviate both the histopathological changes and the physiological effects of urinary dysfunction.

As previously mentioned, the CP1 infection in NOD mice develop symptoms of LUTS similar to that of C57BL/6 mice along with pelvic pain (28). While our current study has not assessed the role of mast cells in the NOD mouse model of LUTS with pain, we can hypothesize that mast cells might play a similarly crucial role in the development of inflammation, fibrosis, and urinary dysfunction in NOD mice albeit the type of immune response (Th1/Th2/Th17 skew) might be altered. It would be interesting to assess mast cell inhibition in the NOD model of LUTS associated with pain as a means to address whether this combination treatment of MCS + H1RA could also be an efficient therapeutic intervention for chronic prostatitis / chronic pelvic pain syndrome (CP/CPPS) patients who have LUTS.

In conclusion, our study demonstrates that mast cells appear to play a critical role in the pathogenesis of voiding dysfunction in an uropathogenic *E. coli* induced mouse model of LUTS. Mast cells have been shown to be present in increased numbers in the prostates of patients with BPH/LUTS. Whether these mast cells are functionally active remains to be determined. The observations from this study show that blockade of mast cell function is therapeutically effective for ameliorating voiding dysfunction. These studies have important translational implications for men diagnosed with BPH associated LUTS.

## ACKNOWLEDGMENTS

We thank Dr. Simon W. Hayward (Department of Surgery, NorthShore University HealthSystem, Evanston, IL, USA) for providing his insights into the project and providing some of the initial human patient prostate sections for characterizing and standardizing the staining in human tissues.

Histology services were provided by the Northwestern University Research Mouse Histology and Phenotyping Laboratory which is supported by NCI P30-CA060553 awarded to the Robert H Lurie Comprehensive Cancer Center”. Flow cytometry services were provided by the Northwestern University Robert H. Lurie Comprehensive Cancer Center Flow Cytometry Core Facility which is supported by Cancer Center Support Grant (NCI CA060553). The model for Figure 10 was Created with BioRender.com.

## GRANTS

This work was supported by the National Institute of Diabetes and Digestive and Kidney (NIDDK) grant R01DK083609 to Praveen Thumbikat and R01DK117906 to Dr. Simon W. Hayward and Praveen Thumbikat. The funders had no role in study design, data collection and analysis, decision to publish, or preparation of manuscript.

## DISCLOSURES

The authors declare no competing financial interests.

## AUTHOR CONTRIBUTIONS

G.P., D.J.M., and P.T. conceived and designed research. G.P., D.J.M., A.J.B-C., and S.F.M. performed experiments. G.P. prepared figures. G.P. and P.T. analyzed data. G.P., A.J.S., and P.T. interpreted results of experiments. G.P. drafted the manuscript. G.P., A.J.S., and P.T. edited, and finalized the manuscript. All authors approved the final version of the manuscript.

## ABBREVIATIONS

BPH: benign prostatic hyperplasia
CrS: cromolyn sodium salt
CeHCl: cetirizine di-hydrochloride
H1RA: histamine 1 receptor antagonist
IHC: Immunohistochemical
IMI: inter-micturition interval
LUTS: lower urinary tract symptoms
MCS: mast cell stabilizer
MLC: myosin light chain
PAR2: protease activated receptor 2
t.u.: transurethral
UPR: urine production rate

## Notes

### Competing Interest Statement

The authors have declared no competing interest.

